# Epithelial N-methyl-D-aspartate (NMDA) receptors mediate renal vasodilation by affecting kidney autoregulation

**DOI:** 10.1101/2023.12.04.569973

**Authors:** Cesar A. Romero, Jasmine Lim, Hong Wang, Brandi M. Wynne, Peipei Ma, Yao Jing, Dennis C. Liotta, Michael D’Erasmo, Stephen F. Traynelis, Douglas C. Eaton, Susan M. Wall

## Abstract

**Background:** N-methyl-D-aspartate receptor (NMDAR) are amino acid receptors that are well studied in brain physiology; however, their role in kidney is poorly understood. Nonetheless, NMDAR inhibitors can increase serum K+ and reduce GFR, which suggests they have an important physiological role in the kidney. We hypothesized that NMDARs in the distal nephron induce afferent-arteriole vasodilation through the vasodilator mechanism connecting-tubule-glomerular feedback (CNTGF) that involves ENaC activation.

**Methods and results:** Using a tubule-specific transcriptome database combined with molecular biology and microscopy techniques, we showed kidney expression of NMDAR subunits along the nephron and specifically in ENaC-positive cells. This receptor is expressed in both male and female mice, with higher abundance in females (p=0.02). Microperfusing NMDAR agonists into the connecting tubule induced afferent-arteriole vasodilation (EC_50_ 10.7 vs. 24.5 mM; p<0.001) that was blunted or eliminated with the use of NMDAR blocker MK-801 or with the ENaC inhibitor Benzamil, indicating a dependence on CNTGF of the NMDAR-induced vasodilation. In vivo, we confirmed this CNTGF-associated vasodilation using kidney micropuncture (Stop-flow pressure 37.9±2.6 vs. 28.6±1.9 mmHg, NMDAR agonist vs vehicle; p<0.01). We explored NMDAR and ENaC channel interaction by using mpkCCD cells and split-open connecting tubules. We observed increased amiloride-sensitive current following NMDAR activation that was prevented by MK-801 (1.14 vs. 0.4 *μ*Amp; p=0.03). In split-open tubules, NMDAR activation increased ENaC activity (Npo Vehicle vs. NMDA; p=0.04).

**Conclusion:** NMDARs are expressed along the nephron, including ENaC-positive cells, with higher expression in females. Epithelial NMDAR mediates renal vasodilation through the connecting-tubule-glomerular feedback, by increasing ENaC activity.

## Introduction

N-methyl-D-aspartate receptor (NMDAR) inhibitors are one of the first classes of drugs available for treatment of early state dementia. It is estimated that they make up 1% of all prescriptions, and 20% of patients with cognitive impairment receive them daily.(7, 16) In humans, the use of NMDAR antagonists has been linked to hyperkalemia and creatinine increase without a clear mechanism.(14) Similarly, in rats, NMDAR antagonists decreased glomerular filtration rate (GFR) and cortical plasma flow, and reduced potassium excretion.(8) NMDAR are Ca2+-permeable ligand-gated cation-selective channels that bind amino acids such as glutamate and glycine, as well as other naturally occurring substances such as quinolinic acid.(6) NMDAR are heterotretamers consisting of two mandatory GluN1 subunits in combination with two other subunits that can be a combination of GluN2A, GluN2B, GluN2C, GluN2D, GluN3A or GluN3B.(6)

In the kidney, NMDAR expression has been reported in podocytes, proximal tubules, and the medulla region.(5, 6) In a previous report, NMDARs were linked to an increase in renal blood flow and GFR that was not mediated by the renal nerves. Deng and Thompson (2009) showed that glycine infusion inside the tubule increased single nephron GFR independent of tubule-glomerular feedback (TGF), but the mechanism was unknown.(4)

More recently a new vasodilator feedback mechanism has been described in the aldosterone-sensitive distal nephron, called connecting-tubule-glomerular feedback (CNTGF).(12) CNTGF is an autoregulatory mechanism that antagonizes and resets vasoconstrictor tubule-glomerular feedback (TGF) by inducing afferent arteriole vasodilation.(15) When high sodium is detected by the epithelial sodium channel (ENaC) in the connecting tubule, the principal cells release prostaglandins and epoxy-acids from the basolateral part of the tubule. These act in a paracrine fashion in the attached portion of the afferent arteriole, inducing afferent-arteriole vasodilation, and increasing glomerular pressure and GFR.(10) ENaC is the only described activator of CNTGF, and also ENaC is participating in K+ homeostasis, inducing K+ secretion by generating a lumen negative potential in the distal nephron. Thus, we hypothesize that NMDAR activation in the kidney is inducing renal vasodilation through the activation of ENaC and consequently initiating CNTGF . Thus, the use of NMDA inhibitors may decrease vasodilation and potassium excretion.

These data provide a novel understanding of the effects of NMDAR and NMDAR inhibitors on kidney autoregulation and tubular transport.

## Methods

### Kidney collection

All protocols followed the National Institutes of Health Guide for the Care and Use of Laboratory Animals and were approved by Emory University and Henry Ford Hospital Institutional Animal Care and Use Committee. Rodents were anesthetized with inhalation isoflurane (4% induction and 2% maintenance). The abdomen and chest were opened and organs exposed. The right atrium was excised and immediately perfused through the left ventricle with cold heparinized saline (20 IU/ml) at 0.5ml/min, until the fluid exiting the right atrium was clear (∼ 5 min). One of the kidneys was harvested for molecular biology analysis and the remaining kidney was perfused with formalin 4% in PBS for 10-15 minutes for fixation.

### RNA expression in epithelial cells

To explore NMDAR RNA expression along the different tubule segments of the nephron, we used the NHI web tool resource which provides full-length RNA-sequence transcriptomes of 14 microdissected segments of mouse kidney tubules.(2) The online tool (https://esbl.nhlbi.nih.gov/MRECA/Nephron/) has been validated and compared with previous databases and protein expression. We explored RNA transcription on each different tubule subsegment of *Grin1, Grin2A, Grin2B, Grin2C, Grin2D, Grin3A* and *Grin3B* genes for each of the NMDAR subunits. RNA sequences read are expressed as logarithmic transformed transcripts per million (TPM).

### Immunoblot

Whole kidney lysates were generated by homogenization using an Omni THQ tissue homogenizer, in lysis solution containing Roche Complete Protease Inhibitor Cocktail (Cat# 04693159001), and then centrifuged at 1,000 g for 15 minutes at 4°C. To enable equal protein loading in each lane, protein content in the soluble fraction of homogenates was measured using a RC-PC protein assay kit (DC Protein Assay Kit, Bio-Rad, Hercules, CA), and then dissolved in Laemmli buffer.

Lysate protein were separated by mini SDS-PAGE on 4.5-12% acrylamide gels, electroblotted to PVDF membranes (Immobilon, Millipore, Bedford, MA), blocked with Odyssey blocking buffer (LI-COR Biosciences) following the manufacturer’s instructions and then incubated with a primary antibody (anti-NMDAR type 1; 1:1,000; StressMarq, S308-48) overnight at 4°C, followed by incubation for 2 h at room temperature with Alexa Fluor 680-linked anti-mouse IgG (A21058; Invitrogen, 1:5000). Signals were visualized with an Odyssey Infrared Imaging System (LI-COR Biosciences). Band density was quantified using *Image Studio*^*tm*^ acquisition software version 5.2 (LI-COR Biosciences). For each sample, band density was normalized to actin (A2066; Sigma-Aldrich, 1:1,000).

### Immunostaining

Paraffin-embedded kidney sections were deparaffinized and antigen retrieval was performed using Trilogy solution (Cell Marque™, 920P). Tissue incubation with antibodies against GluN1 (StressMarq, S308-48) at 1:5000 dilution was performed overnight at 4°C. Then, the sections were incubated with a secondary biotinylated antibody for 30 minutes at room temperature, using a horse anti-mouse antibody attached to Avidin/Biotin Systems, Mouse on Mouse (M.O.M.) Systems Immunodetection Kit (Vector, BMK-2202). Hematoxylin nucleus counterstaining was performed. A negative control slide without primary antibody incubation was included in all experiments, as well as mouse brain tissue as a positive control.

Frozen kidney sections were fixed with acetone and immunostained with an antibody against GluN1 (StressMarq, S308-48, at 1:50 dilution). A mouse anti-mouse antibody conjugated with Alexa Fluor 647 was used (Invitrogen A21058). 4′,6′-Diamidino-2-phenylindole staining was used to counterstain cellular nuclei. Double-staining for the GluN1 and the principal cell markers of the epithelial sodium channel (ENaC) was performed on 6-*μ*m kidney frozen sections using anti-alpha subunit ENaC channel (rabbit anti-mouse, StressMarq, SPC-403). A goat anti-rabbit IgG secondary antibody conjugated to Alexa Fluor 488 (Ab150077 Abcam) was used.

### Tubule microperfusion

We performed a double tubule-vascular microperfusion of a microdissected CNT with the attached afferent arterioles from rabbit kidney as described before.(11, 13) Briefly, New Zealand White (NZW) male rabbits weighing 1.5–2 kg (Myrtle’s Rabbitry) were given standard chow (Ralston Purina, St. Louis, MO) and tap water ad libitum and anesthetized with ketamine (50 mg/kg), xylazine (10 mg/kg), and pentobarbital sodium (25 mg/kg iv). All protocols followed the National Institutes of Health Guide for the Care and Use of Laboratory Animals and were approved by Henry Ford Health System’s Institutional Animal Care and Use Committee. We used rabbits because their CNTs are well-demarcated and microdissection of the CNT and attached afferent arteriole (Af-Art) is feasible in comparison with rats or mice. A single superficial Af-Art with its intact glomerulus was dissected along with the adherent CNT. The microdissected complex was transferred to a temperature-regulated perfusion chamber mounted on an inverted microscope. Both the Af-Art and CNT were cannulated and perfused with an array of concentric glass pipettes as described previously.(1, 12) A small exchange pipette inserted inside the perfusion pipette allows us to exchange all of the fluid inside the perfusion line in a few seconds, while keeping the holding and perfusion pipettes in place. The CNT was perfused by a microperfusion pump (Harvard Apparatus, Holliston, MA) and set to 20 nl/min, which is within the range of physiological flow rates. The afferent arteriole was perfused with modified essential medium (MEM) containing 5% BSA, gassed with room air. Intraluminal pressure was measured by Landis’ technique and maintained at 60 mm Hg. The CNT perfusion solution contained (in mmol/l) 4 KHCO_3_, 10 HEPES, 0.5 Na acetate, 0.5 Na lactate, 0.5 K_2_HPO_4_, 1.2 MgSO_4_, 1 CaCO_3_, and 5.5 glucose, and the final NaCl concentration was 0, 5, 10, 20, 40, and 80 mol/l, respectively. Afferent arterioles were preconstricted with norepinephrine (NE; 2–5 × 10−7 mol/l). Two consecutive concentration-response curves were generated by increasing the CNT luminal NaCl from 0 to 5, 10, 20, 40, and 80 mM (each NaCl concentration was maintained for 5 minutes) with or without a mix of glutamate (1 mM) and glycine (1 mM), and combining it with the non-selective NMDAR inhibitor MK-801 (0.1 mM). Benzamil (1 *μ*M) was used in the perfusate to evaluate the presence of CNTGF-induced vasodilation. The diameter of afferent arterioles was measured in the region of maximal response to NE at three sites, 3–5 μm apart, and expressed as the average of these three measurements. All drugs were added to the CNT lumen. All chemicals were purchased from Sigma (St. Louis, MO).

### Stop-Flow pressure

Sprague-Dawley rats, ∼300 gr body weight, fed on normal salt diet, were used for these experiments. After anesthesia (thiobutabarbital 125 mg/kg, ip), a tracheostomy was performed. Body temperature was maintained at 37.5°C using a feedback-controlled, heated surgical table. The jugular vein was catheterized for continuous infusion of 0.9% NaCl at 1.5 ml/hr. The femoral artery was also catheterized with PE-10 tubing attached to a pressure transducer (AD Instruments) to continuously monitor blood pressure. After a flank incision, the kidney was exposed and placed in a Lucite cup, surrounded with saline-soaked cotton. The bladder was also catheterized to freely excrete urine. After a 30-to 45-minute equilibration period, proximal tubules were identified by injecting a phosphate buffer (pH 7.4) containing 7% of a green dye (made from FD&C yellow no. 5, FD&C yellow no. 6 and FD&C blue no. 1; Crompton & Knowles, Middlebury, CT) in a random proximal segment using a glass pipette (OD 6–8 μm at the tip). A nephron was used when at least three downstream proximal segments were identified, indicating an early proximal puncture site. An early proximal tubule was blocked with grease (Type T Medium Temperature Vacuum Grease; Apiezon). A perfusion pipette (OD 8–10 μm) attached to a nanoliter infusion pump was inserted into the last superficial proximal segment to perfuse the loop of Henle and distal nephron, with a solution containing (in mmol/l): 140 NaCl, 10 HEPES, 1 CaCO_3_, 0.5 K_2_HPO_4_, 4 KHCO_3_, 1.2 MgSO_4_, 5.5 glucose, 0.5 Na acetate, 0.5 Na lactate, and 0.5% of the above-mentioned green dye (pH 7.4). To measure stop-flow pressure (PSF), a pressure pipette (OD 3–4 μm) attached to a micropressure system (model 900A; World Precision Instruments, Sarasota, FL) was inserted into an early proximal segment. The late proximal perfusion rate was increased from 0 to 10, 20, 30, and 40 nl/min while PSF was measured. Each perfusion rate was maintained for 75 seconds, and the measurement range of the PSF response was 45 to 75 seconds. We performed two consecutive tests of PSF response in each experiment while adding glycine plus glutamate (1 mM) with or without ENaC inhibitor benzamil (10^−6^ M). CNTGF was measured by examining the PSF difference when NMDAR agonist was infused, with and without Benzamil.

### ENaC activity

mpkCCD cells, an immortal cell line derived from cortical collecting duct transgenic mice, were maintained in culture as described before.(9) Cells were grown on DMEM/Hams F12 with 2% fetal bovine serum and standard cell culture hormones (Dexamethasone 1 1 mM; mM, T3 1 *μ*M, ITS 100X Gibco cat # 41400045) on 6-well, 4-μm-pore-size Transwell polycarbonate membranes (Costar, Cambridge, MA). All cells were maintained in medium at 37°C, 5% CO_2_. After monolayers exhibited appropriate transepithelial resistance, the media were aspirated and the cells were washed twice with phosphate-buffered saline. Fully defined media, without FBS or hormones, were then applied to both sides of the monolayer. After 48 hours of incubation in charcoal-stripped media, the experiment was begun. Cells were then treated with vehicle (media), N-methyl-D-aspartate (100 *μ*M, NMDAR agonist) or MK-801 (0.1 mM, NMDAR antagonist). The NMDAR 2C/D selective inhibitor 997-74 (1 *μ*M) was also evaluated.(3) Cell monolayer transepithelial voltage (VTE) and resistance (RTE) were measured using an epithelial volt-ohmmeter equipped with stick electrodes (World Precision Instruments, Sarasota, FL). The equivalent transepithelial current (ITE) was calculated according to Ohm’s law (ITE = VTE/RTE) and then corrected for the Transwell insert surface area.

#### Single-Channel Recordings in split-open tubules

Microdissected CNT/cortical collection ducts were used in all patch clamp studies. Micropipettes were pulled from filamented borosilicate glass capillaries (TW-150F; World Precision Instruments) with a two-stage vertical puller (Narishige, Tokyo, Japan). The resistances of the pipettes were between 7 and 10 MΩ when filled with and immersed in patch solution containing the following (in mM): 96 NaCl, 3.4 KCl, 0.8 MgCl_2_, 0.8 CaCl2, 10 HEPES, with pH adjusted to 7.4 by NaOH. Single-channel recordings were made for approximately 8 to 10 minutes at pipette holding potentials of 0 or 20 mV with and without the presence of NMDA (100 *μ*M). Channel currents were recorded at 1 kHz with an Axopatch 1-D amplifier (Molecular Devices Inc., Sunnyvale, CA) with a low-pass 100-Hz 8-pole Bessel filter. Channel activity per patch was determined during an 8-to 10-minute recording period. As a measure of epithelial sodium channel activity, NPo was determined using pCLAMP 10 software (Molecular Devices Inc.). The channel NPo can be calculated as the product of the total number of channels (*N*) in a patch or the open probability of a single channel (*P*_o_). The channel density per patch, N, presented in this paper, is defined as the maximum number of unitary current transitions during 8 to 10 minutes of single-channel recording. This was calculated for all patches including those patches without any observable active channels. The open probability of a single channel (Po) can be calculated by dividing NPo by the number of channels in a patch. Po was calculated only for patches with active channels because we cannot determine if the complete absence of channel activity is due to the absence of channels or the presence of channels with zero open probability.

## Results

### NMDAR expression in the Kidney

The *Grin1* subunit gene transcript and GluN1 protein expression were explored in the kidney. Using an NIH web-based resource, we explored the transcripts for mandatory subunit *Grin1* and other NMDAR subunits from microdissected tubules (Figure 1). *Grin1* transcript expression was found in nearly all nephron segments except the s2-s3 segment of the proximal tubule and the medullary portion of the thick ascending limb of the loop of Henle. Stronger *Grin1* transcription compared to the cortex was observed in the outer medullary region of the nephron (loop of Henle segments and OMCD). In the cortex, in the first segments of the proximal tubules (PTS1), most NMDA receptor subunits are expressed except for *Grin2C*, which is relatively abundant in DCT. In the CNT and CCD, in addition to the mandatory GluN1 subunits, *Grin2D* and trace amounts of *Grin3A* and *Grin3B* are expressed (Figure 1A).

**Figure 1.**
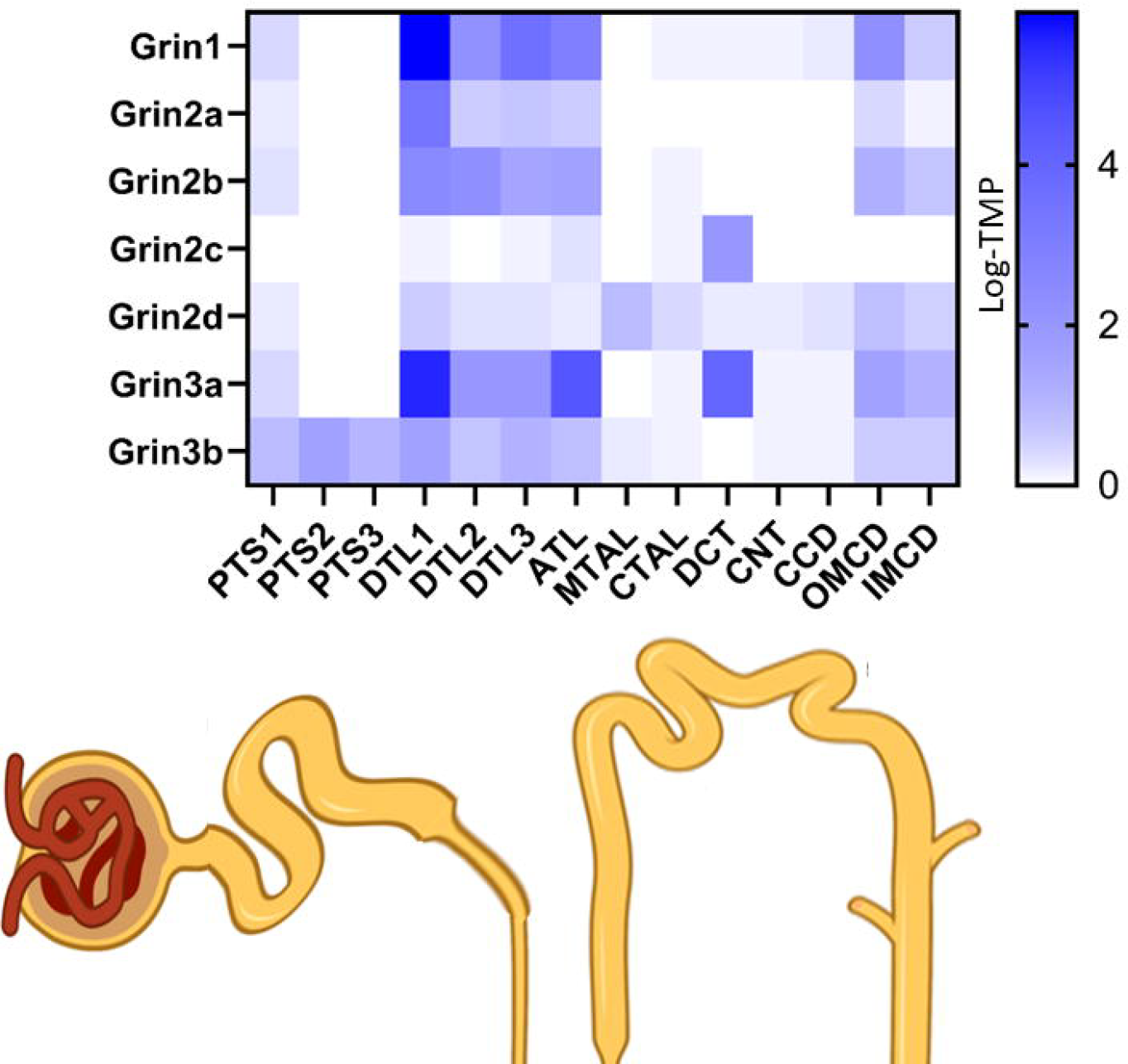
NMDAR RNA expression along the nephron. The mandatory Grin 1 subunit is expressed in all the renal segments, especially in the medulla regions of the nephron. In the medulla region all NMDAR subunit are expressed, while in the cortex in addition to Grin1, Grin2D is the majority expressed in the distal nephron in addition to grin2C and Grin3A in DCT and in less magnitude in CNT and CCD.TMP: Transcript per millon. [PTS1] initial segment of proximal tubule; [PTS2] straight segment of proximal tubule; [PTS3] last segment of the proximal straight tubule in the outer stripe of outer medulla; [DTL1] short descending limb of the loop of Henle; [DTL2] long descending limb of the loop of Henle in the outer medulla; [DTL3] long descending limb of the loop of Henle in the inner medulla; [ATL]) thin ascending limb of the loop of Henle; [MTAL] medullary thick ascending limb of the loop of Henle; [CTAL] cortical thick ascending limb of the loop of Henle; [DCT] distal convoluted tubule; [CNT] connecting tubule, cortical collecting duct [CCD], outer medullary collecting duct [OMCD], and inner medullary collecting duct [IMCD])

GluN1 protein labeling was detected in both the cortical and medullary regions, with preponderant expression in the outer medulla, in line with the RNA transcription pattern (Figure 2A-B). Expression of GluN1 was observed along the tubules and in the glomerulus (Figure 2C). GluN1 localized to both the luminal and basolateral regions of these tubules, as well as to the cytoplasm (Arrows Figure 2C). Using ENaC as a marker of aldosterone sensitive segment, we confirmed the protein expression of GluN1 in these regions. (Figure 2D-F). Comparing males and females, we observed a significant difference in NMDAR expression (Figure 3A), where females exhibit an average fourfold increase in renal NMDAR expression (male vs. female 1.4±0.8 vs 4.9±2.6, p=0.02) (Figure 3B).

**Figure 2.**
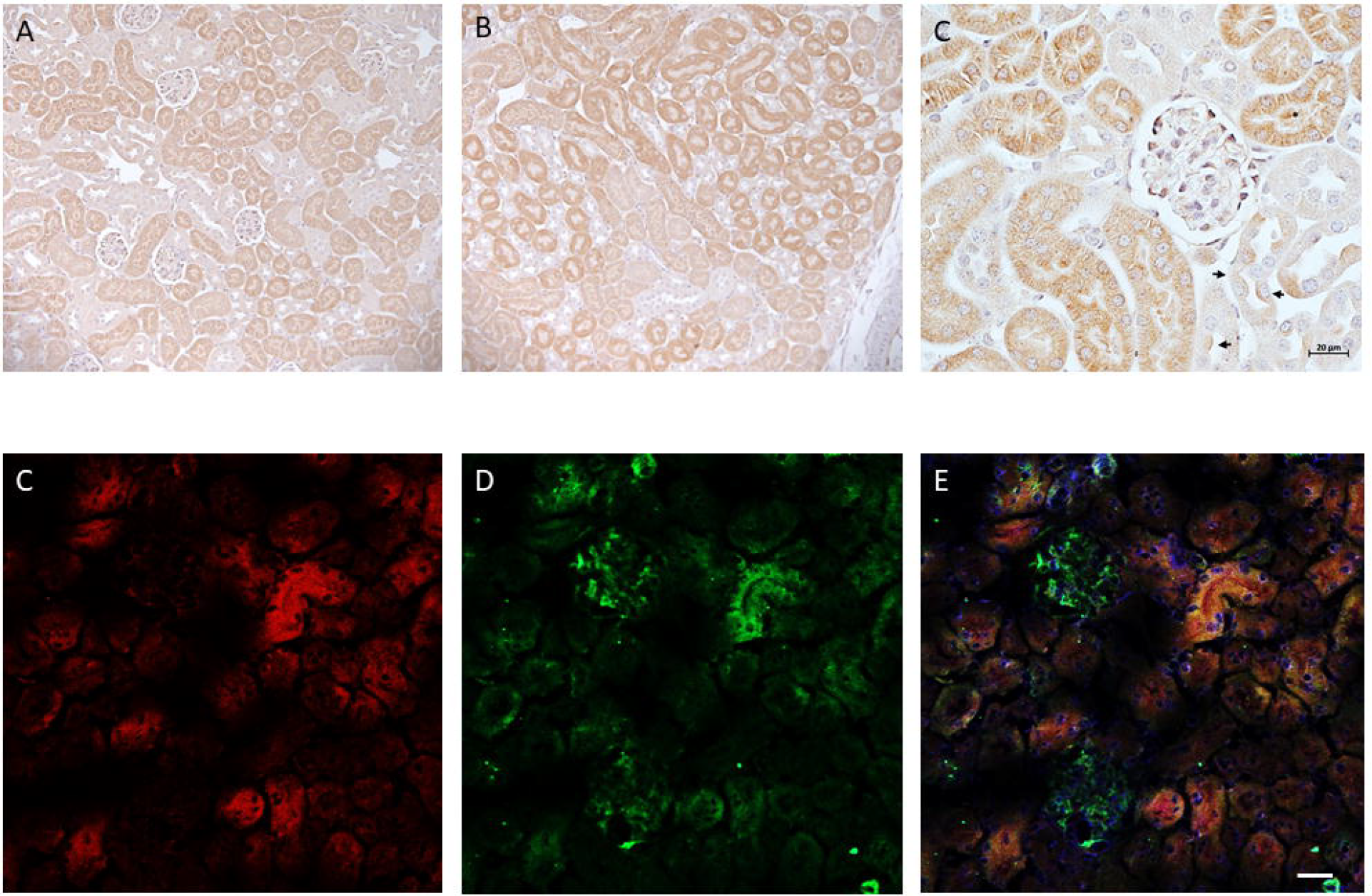
Kidney expression of NMDAR type 1 (GlutN1). GlutN1 is expressed in both cortical (Figure 2A) and medullary region of the kidney (Figure 2B) where strong signal was detected. In the cortex, GlutN1 is expressed several nephron segments and glomerulus (Figure 2C). At higher magnification, GlutN1 disposicition is even in the cytoplasm with perinuclear reinforcement in some cells (star) and while in other segments shows an apical expression (arrows). Figure 2C and 2D shows GlutN1 expression (red) and ENaC alpha subunit (Green) in cortex. Figure 2E shows the co-expression of both protein (yellow) in dotal nephron. Scale bar= 20 *μ*m.

**Figure 3.**
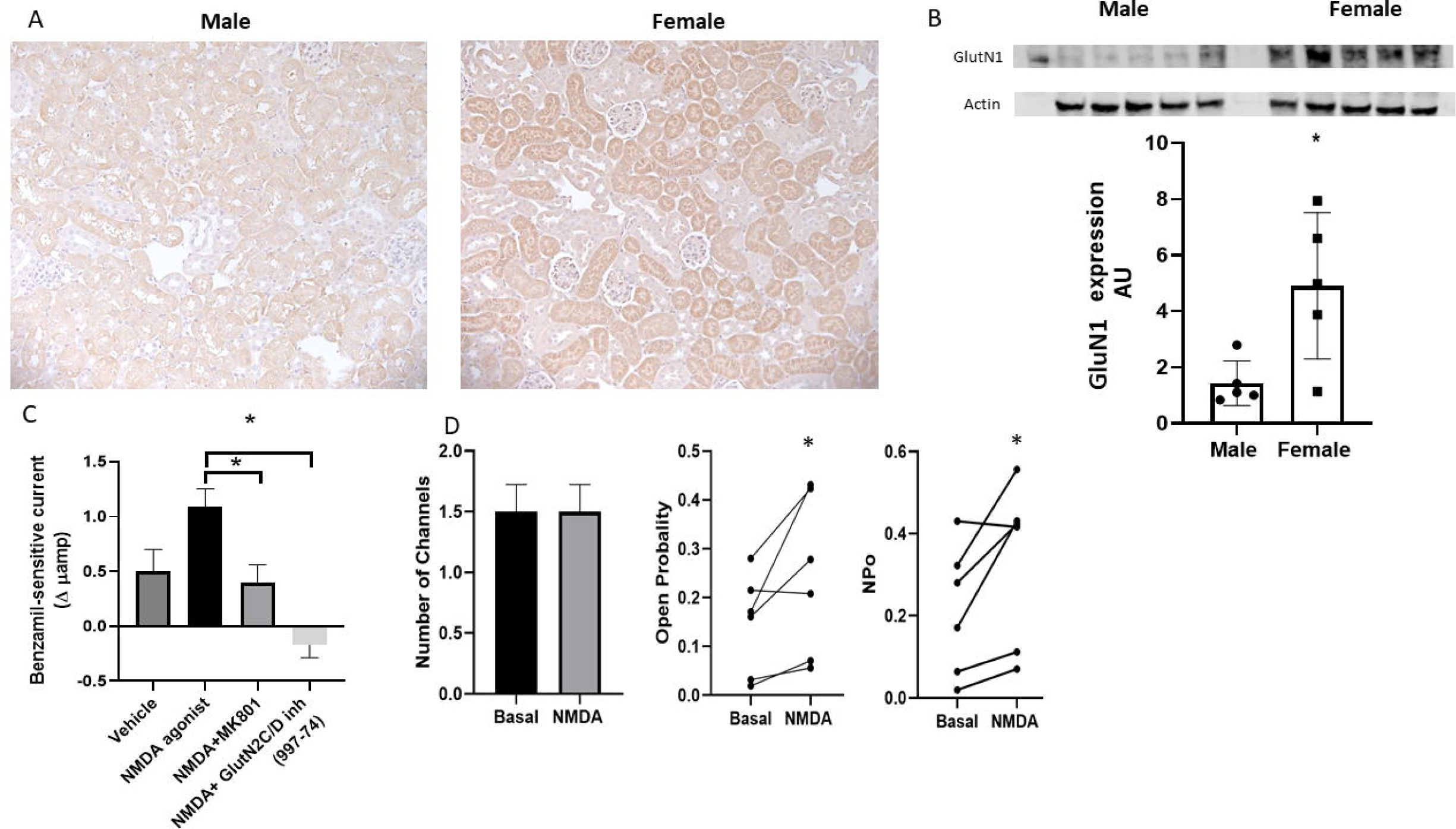
Sex difference in NMDAR type 1 and its interaction with ENaC. Figure 3A compare the expression of GlutN1 in cortex of male and female mice. Male mice exibit a weak staining in comparison to female. Figure 3B shows total kidney expression quantification between male and female. Figure 3 C Figure 3C shows benzamil-sensitive transepithelial current using mpkCCD cells under the presence of NMDAR agonist, MK801 (non-selective blocker) and a selective GLutN2C/D inhibitor (997-74) The presence of NMDA increase the transepithelial current, which is blunted when the non-selective blocker is used and totally abolished whith the use of the selective inhibitor. Figure 2D shows an increase ENaC activity (NPo) at exprense of the open probability of the channel with not changes in the amount functional channels. AU: Arbitrary units (GlutN1/Actine ratio). Δ trasnepithelial current difference. *p<0.05

### NMDAR activation induces CNTGF activation

To determine if the NMDARs in CNT induce renal vasodilation and if that vasodilation occurs through activation of CNTGF, we performed double tubule-vascular microperfusion experiments in microdissected CNT with the attached afferent arteriole. This approach allowed us to selectively evaluate the effect of epithelial NMDAR activation over afferent-arteriole tone. Figure 4A shows the classical afferent-arteriole vasodilation observed after increasing sodium chloride concentration in the CNT. When a mix of glutamate and glycine (agonist and co-agonist) were included in the perfusate, we observed a significant increase of afferent-arteriole vasodilation, shifting the curve to the left. Thus the amount of sodium chloride necessary to induce half of the maximal vasodilaton was dramatically reduced in the presence of NMDAr agonist (EC50 10.7 vs. 24.5 mM; p<0.001). To confirm that the vasodilation observed was related to NMDAR activation, we explored this response in the presence of the NMDAR blocker MK-801, which blocks NMDARs regardless of subunit composition. As shown in Figure 4B, the use of MK-801 shifted the curve to the right, indicating that the increased vasodilation was due to NMDAR activation. To confirm that this vasodilation was ENaC-dependent and related to the autoregulatory mechanism CNTGF, we added the CNTGF-blocker benzamil (ENaC inhibitor). Figure 4C shows that the co-infusion of amino acids and benzamil totally blocked the amino acid-induced afferent-arteriole vasodilation response, indicating that the vasodilation induced by activation of NMDAR was totally dependent on CNTGF. Figure 4D shows that in the absence of luminal glutamate and glycine, MK-801 has no effect. To evaluate the intratubular effect of glutamate/glycine on renal vasodilation in vivo, we measured CNTGF in rats using micropuncture experiments and evaluating stop-flow pressure (PSF), a surrogate for glomerular pressure. Thus, dilation of afferent arterioles induce an increase in glomerular pressure that is transmitted to the tubule where the luminal flow was stopped, acting as a chamber. Figure 4E shows the changes in stop-flow pressure after increasing flow of vehicle (saline). The physiological response to an increased flow in the distal nephron is a decrease in stop-flow pressure. This decreasing curve is a representation of the combined TGF and CNTGF response where TGF predominate inducing a vasoconstriction. When the same experiment is performed with a mix of amino acids, SFP is almost without change, indicating a reduced vasoconstriction response. Figure 4F, demonstrate that the diminished vasoconstrictive response is due to an exaggerated CNTGF activation, because when benzamil (CNTGF blocker) is infused, the vasoconstrictive response due to TGF is unmasked. Figure 4G shows the CNTGF activity in the absence of amino acids. By comparing the CNTGF magnitude with amino acid infusion versus Vehicle, we observed that CNTGF is significantly enhanced (Figure 4H). These data indicate that luminal amino acids glutamate and glycine activate NMDAR, and NMDAR induces vasodilation and increased glomerular pressure through CNTGF.

**Figure 4.**
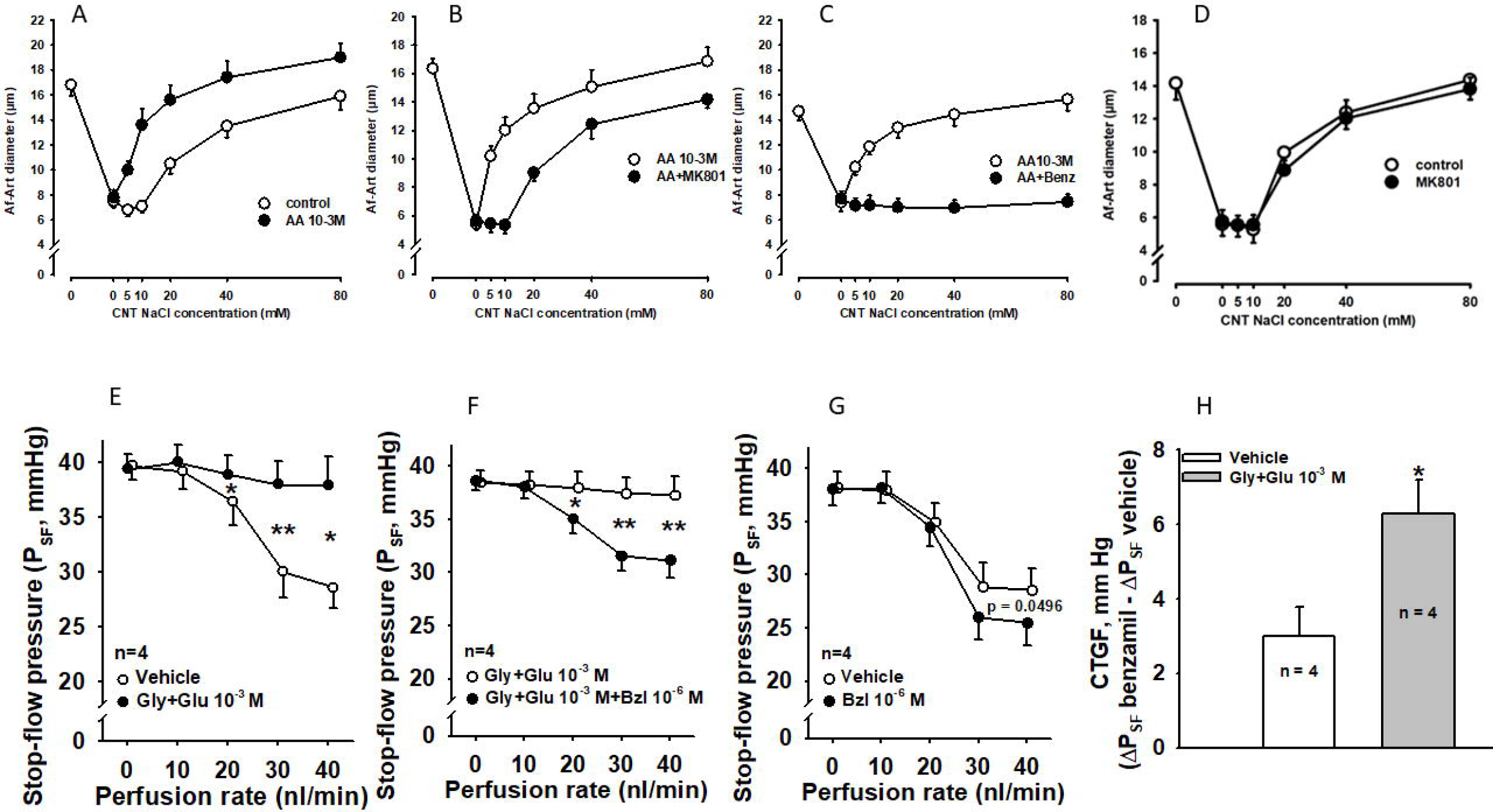
NMDAR induce afferent arteriole vasodilation. **Panel A-D** represent the afferent arteriole (Af-Art) diameter change after the intratubular infusion of increasing sodium chloride concentration combined or not with different agonist and pharmacological inhibitors in the connecting tubule (CNT) (open circles). Co-infusion of sodium chloride with glutamate and glycine increase the Af-art vasodilation, by shifting the curve to the left (closed circle). Figure 4B demonstrate that the addition of the non-selective NMDAR blocker MK801 restore the curve to the right (closed circles). Figure 4C shows that the co-infusion of the ENaC blocker benzamil totally avoid the Af-Art vasodilation in the presence of amino acids, indicating the total dependence on ENaC and CNTGF for the amino acid induced vasodilation (closed circles). Figure 4D shows no effect of MK801 on Af-Art vasodilation in absence of amino acids. **Panel E-H** represent the Stop-flow pressure (SFP) results in-vivo during micropuncture. Figure 4E shows a decrease in SFP, a surrogate of Af-Art constriction after increasing distal flow of sodium chloride. When a mix of amino acids is added to the perfusate, SFP is almost constant, indicating a reduced vasoconstriction response. Figure 4F, demonstrate that the diminished vasoconstrictive response is due to an exaggerated CNTGF activation, because when benzamil (CNTGF blocker) is infused, the vasoconstrictive response due to TGF is unmasked (closed circles). Figure 4G shows the CNTGF activity in the absence of amino acids. By comparing the CNTGF magnitude with amino acid infusion versus vehicle, we observed that CNTGF is significantly enhanced (Figure 4H).

### NMDAR enhance ENaC activity

The micropuncture and microperfusion experiments indicate that NMDAR-induced vasodilation is dependent on CNTGF. Since ENaC stimulates CNTGF, we explore the relationship between ENaC and NMDAR. To do so, we evaluated the localization of ENaC and GluN1 using confocal microscopy (Figure 2D-F). Figures 2D and 2E show the GluN1 (red) and alpha subunit (green) of ENaC. Figure 4F shows the colocalization of both proteins (yellow). This colocalization is observed in several subcellular compartments, especially in the apical region of the cell. We conclude that GluN1 and ENaC colocalize to cells in the aldosterone-sensitive region of the nephron.

We then evaluated the effect of NMDAR agonists and antagonists on benzamil-sensitive transepithelial current in mpkCCD cells, a cell model of kidney principal cells (Figure 3C). To do so, benzamil-sensitive current was measured in mpkCCD cells treated with either the agonist NMDA or vehicle. As shown, benzamil-sensitive current increased with the application of NMDA. However, this response was prevented when cells were co-incubated with the NMDAR blocker MK-801 or with the GluN2C/D-selective negative allosteric modulator, 997-74.(3)

To confirm our observation that NMDAR activation increases ENaC activity, we performed single channel recordings of principal cell apical membrane in split-open mouse CCDs that were incubated with either NMDA or vehicle (Fig. 3D). We observed that while NMDA did not affect ENaC channel surface density (N), it significantly increased ENaC-open probability (Po), which led to increased ENaC channel activity (NPo). We conclude that principal cell NMDAR activation increases ENaC activity.

## Discussion

Our main findings indicate that NMDAR participates in kidney autoregulation by inducing afferent-arteriole vasodilation through the mechanism CNTGF. The CNTGF activation is mediated by an increase in ENaC activity induced by NMDAR.

These findings have important implications for understanding how NMDAR inhibitors used in humans with cognitive impairment may induce renal and metabolic changes, such as GFR reduction and hyperkalemia. Previous studies have shown that CNTGF is highly activated by high-salt diets and when nephron mass is reduced. Additionally, the ENaC generates an electric gradient which induces potassium secretion through the ROMK channel. Inhibition of ENaC is a well-known cause of potassium-sparing and hyperkalemia in some cases.

The effect of amino acids in kidney blood flow has long been known, and best shown by being the basis of the renal functional reserve test. However, the mechanism behind this renal functional vasodilation has been controversial. Our data may indicate that CNT participate in afferent arteriole vasodilation induced by amino-acids and most of the vasodilation effect is related to NMDAR and the CNTGF mechanism.

Our *in vitro* approach, with tubular microperfusion, allowed us to determine that the NMDAR effect was induced in the luminal side of epithelial cells, specific to the connecting tubule, with no interaction of NMDAR expressed in the glomerulus or the afferent arteriole.

Here, we propose a novel interaction between NMDAR and ENaC activity, which poses a completely new regulatory mechanism for not only CNTGF but for ENaC. This opens a new chapter in ENaC regulation, tubular transport, and sodium and potassium homeostasis. Thus, with the presented data we highlight a novel role of ENaC, in NaCl and K+ balance, not only just by mediating NaCl absorption, which increases the driving force for K secretion, but rather, by changing renal blood flow.

We also show a clear difference between the sexes in renal NMDAR expression. Future research is needed to explore the mechanism associated with this sex difference and to evaluate the differential impact of the mechanism on renal disease in males and females.

Unfortunately, this study was not able to explore which specific NMDAR subtypes are mediating these effects. Future research that utilizes additional subunit-selective NMDAR inhibitors and genetically-modified animals will be necessary to determine the subunit composition of the NMDARs mediating this effect. However, our *in vitro* experiments, in addition to the RNA transcriptome data, suggest that NMDAR 2D and/or 2C subunits are the main candidates for the vaso-regulatory actions of NMDARs. Additionally, the RNA transcriptome database used in this study did not compare expression in male and female animals. Thus, sex-related differences in expression of different RNA subunits needs to be confirmed in future studies.

We conclude that NMDAR are expressed along the mouse nephron, including ENaC-positive cells, with higher expression in females. Epithelial NMDAR mediates renal vasodilation by activating CNTGF. The activation of CNTGF is mediated by increased ENaC activity after NMDAR activation.

## Authors Contributions

C.A Romero, DC Eaton and SM Wall conceptualized the study and were responsible for funding acquisition, methodology, project administration, resources and supervision. CA Romero, J Lim, H Wang, B Wynne were responsible for data acquisition, analysis and interpretation. P Ma, J Yao, M D’Erasmo, DC Liotta synthesized and supplied 997-74. SF Traynelis, D Liotta, were responsible in the analysis and interpretation of data obtained with the NMDA inhibitors. C.A Romero wrote the original manuscript. All authors reviewed and edited the manuscript.

## Acknowledgments

We thank Oishi Paul and Auriel Moselei for the technical assistance with the mpkCCD cells.

## Data Sharing Statement

All data used in this study are available on reasonable request to the authors.

